# *Leptospira* spp. seroprevalence in humans involved in a cross-sectional study in Garissa and Tana River Counties, Kenya

**DOI:** 10.1101/2020.07.17.208363

**Authors:** Arithi Mutembei, Festus K. Mutai, Damaris Mwololo, John Muriuki, Mark Obonyo, S.W. Kairu-Wanyoike, M. Wainaina, J. Lindahl, E. Ontiri, S. Bukachi, I. Njeru, J. Karanja, R. sang, D. Grace, B. Bett

## Abstract

**Introduction:** Leptospirosis is a neglected bacterial zoonotic infection caused by spirochetes of *Leptospira genus.* Humans get infected through direct or indirect contact with urine of infected animals or environment. It accounts for more than 300,000 severe cases annually worldwide with case fatality rates of over 30%. Costs of diagnosis and treatment for human and animals, disruption of international trade of animals and products, reduced productivity and reproductivity in animals constitute economic importance. In Kenya, leptospirosis burden is significant but under-diagnosis and under-reporting affects the awareness of the disease. This study aimed to determine and compare the sero-prevalence and factors associated with *Leptospira* spp. in the two counties.

**Methods:** We conducted a cross-sectional study that involved apparently healthy people of at least 5 years of age in randomly selected households in Garissa and Tana River Counties. Blood samples were collected and tested for *Leptospira* spp antibodies using IgM ELISA. Standardized structured questionnaires were administered to collect socio-demographic and exposure information. We calculated frequencies and proportions for categorical variables and odds ratios (OR) and 95% confidence interval (CI) to evaluate association between sero-positivity and exposure factors. We used Wilcoxon test to evaluate statistical difference in sero-positivity for continuous variables and calculated test statistic (H) and p-value.

**Results:** A total of 952 subjects were recruited into the study – these included 482 persons from Garissa and 470 from Tana River. The overall sero-prevalence was 26% [(244/952); (CI: 23% to 29%)]. Garissa County had significantly higher *Leptospira* spp. seroprevalence (31%, n = 147; CI: 27% to 35%) compared to Tana River County (21 %, n = 97; CI: 17% to 25%). Being a female (OR=1.6, CI: 1.2-2.2) and engaging in pastoralism (OR=2.7, CI: 1.8-3.9) were significantly associated with higher odds of *Leptospira* spp. seropositivity compared to being a male or working in irrigated areas. The mean altitude of residence of sero-positive patients was 73m ± 21 SD (standard deviation) above sea level and that for sero-negative was 80m ± 22 SD (H=35, p-value = 0.00).

**Conclusion:** This study determined the seroprevalence and risk factors for *Leptospira* spp. exposure in Garissa and Tana River Counties, Kenya. Females in pastoral communities experience high burden of the disease. Enhanced surveillance in humans and animals and further research is required to understand the complex and multifactorial drivers of leptospirosis transmission in the two Counties.

## Introduction

Leptospirosis is a neglected bacterial zoonotic infection caused by spirochetes of the genus *Leptospira.* Humans mainly get infected through direct or indirect contact with urine of infected animals or contaminated environment [1]. *Leptospira* spp. seroprevalence vary significantly between and within countries, based on environmental settings, behavioral risk factors, and socio-demographics. The majority of the morbidities and mortalities occurs in regions which have large subsistence farming, pastoral populations and where the disease is a veterinary health problem [2]. Its impacts are associated with reduced livestock production, high costs of detection and treatment in human and animals, and disruption of international trade of animals and its products.

It is estimated that 7 to 10 million people are infected by leptospirosis globally with about 1.03 million cases and 58,900 deaths occurring due to leptospirosis annually. This translates to approximately 2.90 million Disability Adjusted Life Years [3]. A large proportion of cases (48%) and deaths (42%) occur in adult males within age of 20–49 years [2]. In Africa, *Leptospira* incidence has been estimated to be 95.5 cases per 100,000 and prevalence ranges from 2.3% to 19.8% [4]. A few studies have been done in Kenya on the disease; available data show that in 1987, 7.4% of 353 healthy people in Nyanza region and 16.9% of 130 in the coastal region had *Leptospira* antibodies [5]. Later in 2012, a cross-sectional survey that involved slaughterhouse workers in Busia, Kenya reported a prevalence of 13.4% [6].

Although leptospirosis is known to be one of the high consequence zoonotic disease worldwide, there is limited knowledge on its burden, spatial distribution, and relative importance of the disease in Kenya. The disease has been listed among the top 20 priority diseases in the country [7], but there is inadequate knowledge on its distribution, risk factors and the most vulnerable groups in the country. Its surveillance is inadequate, and there have not been any investments on community sensitization campaigns to improve its detection and reporting. The disease is under-diagnosed and under-reported due to multiple challenges including nonspecific clinical symptoms [8–12]. It is one of the neglected diseases [13,14], and in resource-constrained countries (e.g. Kenya) that has multiple diseases and socio-economic challenges to confront, quantification of leptospirosis burden would guide re-evaluation of disease priority lists [11,15]. This study determined *Leptospira* spp. seroprevalence and risk factors in selected areas in Garissa and Tana River Counties, eastern Kenya.

## Methods

### Study site

The study was conducted in Bura and Hola, Tana River County, and Ijara and Sangailu in Garissa County. These sites have been described earlier [16]. Briefly, Bura site was in an irrigation and settlement scheme that covered 2,100 hectares with an approximate population of more than 2,000 households spread out in 10 villages. Hola study site was in a neighbouring irrigation and settlement scheme that covered 1,011 hectares with 700 farming households in 6 villages. They received approximately 460mm per year which peak in October–December. Their average daily temperatures ranged between 32 - 37°C. Ijara and Sangailu fell under Ijara sub-County which borders Lamu County and Boni forest to the East and Tana River County to the West. This is an arid/semi-arid area where pastoralism is practised. Their annual rainfall ranged between 750mm – 1000 mm while the mean temperature ranged between 15°C - 38°C. Sampling was done from December 2013 to February 2014.

### Study design

The study used a cross-sectional study design with households as the primary sampling unit, and subjects in households as secondary units. In Bura and Hola, lists of households that were used as the sampling frame were obtained from the local irrigation schemes. In Ijara and Sangailu, lists of households were developed with the help of the village headmen.

The number of people/households to sample was determined using a sample size for estimating a population proportion. A minimum sample size of 845 was estimated assuming that *a priori Leptospira* spp. seroprevalence was 50%, level of confidence was 95%, reliability 5% and that the correlation coefficient, *ρ*, of seropositivity outcome at a household level was 0.3. It was further assumed that up to five people per household, *m*, would be sampled.

### Sampling methods Study Population

The sample size was uniformly distributed to all the study areas in order to have 211 subjects or 42 households per site. These households were randomly identified from the sampling frames using computer generated random numbers. Within each household, the household head was asked to identify up to 5 people for sampling. These had to be participants that were healthy above the age of 5 years at the time of study. Children below the age of 5 years were not included in the study.

### Data collection

We administered a standardized questionnaire to the study participants to collect socio-demographic, household and area level data that would be used to determine risk factors for *Leptospira* spp. exposure. Variables collected included age, sex, occupation, education status, total number of people in a household, source of water, geographical coordinates and altitude and the main land use activity per sampling site. These data were collected using the Open Data Kit (ODK) application in android-enabled smartphones. The data were posted at the end of each day to an on-line server at the International Livestock Research Institute (ILRI).

Blood samples were then collected from up to five people in a household. The household was given the authority to identify a household member who would be sampled provided they were more than 5 years. Recruited subjects were seated comfortably for blood collection. Up to 10ml venous blood was drawn from the left median cubital vein after disinfecting the injection site using 70% isopropyl alcohol. A tourniquet was placed about 3-4 inches above the venipuncture site. Sterile butterfly needles and vacutainer tubes were used to draw blood which was collected into non heparinized tubes. Blood samples were transported in a cool box to a field laboratory where they were centrifuged at 3000xg for 10 minutes. Serum harvested was aliquoted to 2ml cryotubes in duplicates and kept in dry ice until transported to ILRI Nairobi where it was preserved at −80⁰C until analyzed.

The samples were screened for anti-*Leptospira* spp. IgM using Panbio^®^ Leptospira IgM ELISA. The kit is known to detect infections caused by a wide range of *L. interrogans* serovars including: hardjo, pomona, copenhageni, australis, madanesis, kremastos, nokolaevo, celledoni, canicola, grippotyphosa, szwajizak, djasiman and tarassov. A case was defined as a positive human serum sample for *Leptospira* antibodies on Panbio® Leptospira IgM ELISA

### Data analysis

Seropositivity was assumed to represent the odds of *Leptospira* exposure in people. Data posted to an online server at ILRI were transferred to a relational database designed using MS Access. These were merged with the laboratory results, cleaned and analysed using descriptive and analytical models.

Descriptive analyses included the determination of the number of subjects, households, and villages sampled. The distribution of each variable was also analysed. *Leptospira* seropositivity was considered as the dependent variable, while gender, age, occupation (pastoralist, farmer, student and other), source of water (borehole, canal, dam and other), livestock ownership (yes/no), family size, land use (pastoral, irrigation, riverine), location (Sangailu, Ijara, Bura and Hola) and altitude were independent variables. Age was initially analysed as a continuous variable, but it was later transformed to categorical variable with three levels – as <20, 20 – 40 and >40 years – when it failed to meet the linearity assumption. Occupations grouped under the other category included businesspersons, employed, driver, village chief and a nurse, among others.

Bivariate analysis involving the outcome and all the categorical variables was performed and *Leptospira* spp. seroprevalences and their 95% confidence intervals were calculated at the various levels of these variables. Chi-square tests were also used to assess crude associations between each factor and the dependent variable (at α = 0.05). The variation in *Leptospira* spp. with continuous variables, including altitude at the sampling site and the size of the household were analysed using T Test. These were succeeded by univariable regression models involving each of the variables using a crude logistic repression model. Variables that were significant or returned a p<0.2 were included in multivariable modelling.

Multivariable analysis used a mixed effects logistic regression model, implemented using *melogit* command in STATA version 14, to account for clustering of data at the household and village levels. A mixture of backward and forward variable selection technique was used involving all the variables that met the criteria specified under the univariable analysis described above. Two random effects variables – village ID and household ID -- were fitted, and their significance assessed using the likelihood ratio test. Fixed and random effects variables were retained in the model if their likelihood ratio tests were significant, assuming a type II error, α, of 0.05. The residual intra-cluster correlation coefficient was estimated using the command *etstat icc* after running the final model. Residual analysis using standardised residuals was conducted to identify outliers.

### Ethical considerations

We sought ethical approval from the African Medical and Research Foundation (AMREF) Ethics and Science Review Committee (ESRC). An approval number provided was AMREF-ESRC P65/2013. Informed consent was sought from participants and assent for children below 18 years. The participants were assured of confidentiality and anonymity. Benefits and/or risks associated with the study were explained to the participants who had freedom to withdraw from the study at any time. The researcher ensured safety of data collected through access restrictions by use of passwords and storage in lockable lockers.

## Results

### Descriptive statistics

A total of 952 subjects were recruited into the study. Their mean age was 29.9 (95% confidence interval: 28.7 – 31.2) years. Most of them were female (59.1%, n = 562). The subjects came from 347 households distributed in 40 villages in Tana River and Garissa counties. The number of subjects sampled per household and village ranged between 1 - 5 and 3-56, respectively. Households however had more members with the mean household size being 8.6 (95% CI: 8.4 – 8.9). The study area had a mean elevation of 78.7 (SE = 0.69) m above sea level A detailed analysis of the characteristics of the subjects is given in Table 1.

**Table 1.**
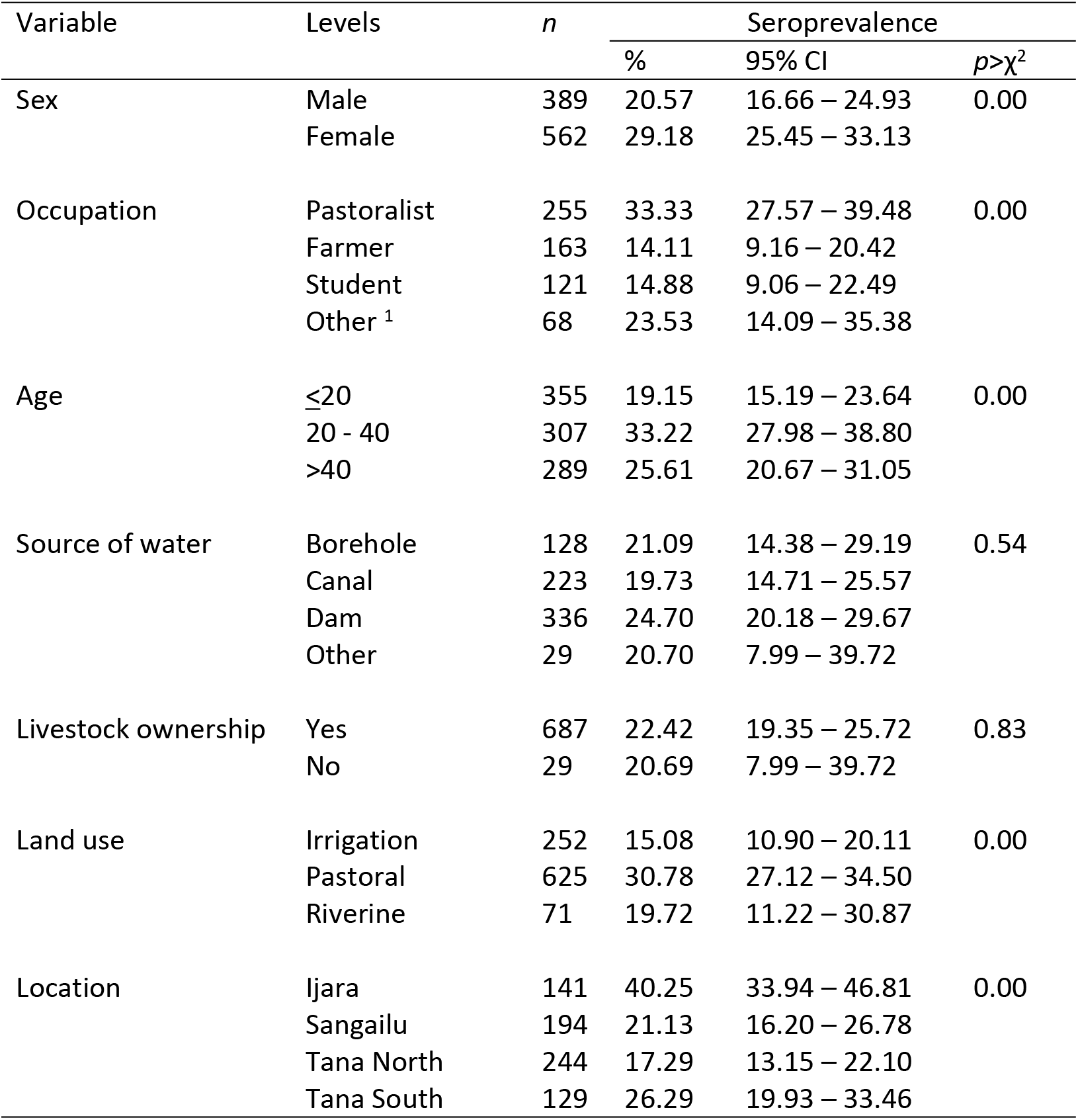
Variation in *Leptospira* spp. seroprevalence by categorical variables defined at the subject, household and area levels

| Variable | Levels | n | Seroprevalence |  |  |
| --- | --- | --- | --- | --- | --- |
| | | | % | 95% CI | $p > \chi^2$ |
| Sex | Male | 389 | 20.57 | 16.66 – 24.93 | 0.00 |
|  | Female | 562 | 29.18 | 25.45 – 33.13 |  |
| Occupation | Pastoralist | 255 | 33.33 | 27.57 – 39.48 | 0.00 |
|  | Farmer | 163 | 14.11 | 9.16 – 20.42 |  |
|  | Student | 121 | 14.88 | 9.06 – 22.49 |  |
|  | Other <sup>1</sup> | 68 | 23.53 | 14.09 – 35.38 |  |
| Age | ≤20 | 355 | 19.15 | 15.19 – 23.64 | 0.00 |
|  | 20 - 40 | 307 | 33.22 | 27.98 – 38.80 |  |
|  | >40 | 289 | 25.61 | 20.67 – 31.05 |  |
| Source of water | Borehole | 128 | 21.09 | 14.38 – 29.19 | 0.54 |
|  | Canal | 223 | 19.73 | 14.71 – 25.57 |  |
|  | Dam | 336 | 24.70 | 20.18 – 29.67 |  |
|  | Other | 29 | 20.70 | 7.99 – 39.72 |  |
| Livestock ownership | Yes | 687 | 22.42 | 19.35 – 25.72 | 0.83 |
|  | No | 29 | 20.69 | 7.99 – 39.72 |  |
| Land use | Irrigation | 252 | 15.08 | 10.90 – 20.11 | 0.00 |
|  | Pastoral | 625 | 30.78 | 27.12 – 34.50 |  |
|  | Riverine | 71 | 19.72 | 11.22 – 30.87 |  |
| Location | Ijara | 141 | 40.25 | 33.94 – 46.81 | 0.00 |
|  | Sangailu | 194 | 21.13 | 16.20 – 26.78 |  |
|  | Tana North | 244 | 17.29 | 13.15 – 22.10 |  |
|  | Tana South | 129 | 26.29 | 19.93 – 33.46 |  |

### *Leptospira* spp. seroprevalence

The overall *Leptospira* spp. seroprevalence was 25.6% (95% CI: 22.9 – 28.5). Table 1 shows the variation in *Leptospira* spp. seroprevalence by categorical variables considered in the study. All the subject-level variables (sex, occupation and age category) had significant effects while the household categorical variables – source of water and ownership of livestock – were not. Area-level variables (land cover and the identify of an area) were also significant.

Two other continuous variables – the size of a household and altitude –were also significantly associated with *Leptospira* spp. seroprevalence. Household sizes for the seropositive subjects were significantly larger (mean of 9.2 persons; 95% CI: 8.7 – 9.7) than the those for seronegative ones (8.4; 8.1 – 8.7). Similarly, the mean elevation for the seropositive subjects (73.0m; 95% CI: 70.3 – 75.7m) was significantly lower than that for seronegative subjects (80.7m; 95% CI: 79.2 – 82.3m).

### Univariable and multivariable analyses

All the variables identified above were fitted in a logit model. All the variables except livestock ownership and source of water were significant. These two insignificant variables were therefore not considered further in the analysis.

The final multivariable model fitted to the data is given in Table 2. Three fixed effects (gender, occupation and location) and one random effect (household) variables met the criterion for inclusion in the model. Age and occupation as well as location and altitude could not be kept in the model at the same time; the presence of occupation in the model made age to be insignificant. Similarly, location rendered altitude insignificant. In both cases, predictors that provided the greatest log-likelihood estimates were preferred.

**Table 2.** Outputs of a random effects logistic regression model used to determine factors that affect the seroprevalence of *Leptospira* spp. in humans in Tana River and Garissa counties, Kenya

| Variable | Level | Odds Ratio |  |  | Z | P>Z |
| --- | --- | --- | --- | --- | --- | --- |
|  |  | Mean | SE | 95% CI |  |  |
| *Fixed effects* |  |  |  |  |  |  |
| Sex | Male | 0.58 | 0.15 | 0.35 - 0.97 | -2.08 | 0.04 |
|  | Female | 1.00 |  |  |  |  |
| Occupation | Farmer | 0.33 | 0.13 | 0.16 - 0.71 | -2.86 | 0.00 |
|  | Other | 0.67 | 0.29 | 0.28 - 1.57 | -0.93 | 0.35 |
|  | Student | 0.36 | 0.14 | 0.17 - 0.77 | -2.63 | 0.01 |
|  | Pastoralist | 1.00 |  |  |  |  |
| Location | Ijara | 3.30 | 1.57 | 1.30 - 8.37 | 2.52 | 0.01 |
|  | Sangailu | 0.75 | 0.33 | 0.32 - 1.76 | -0.65 | 0.51 |
|  | Hola | 1.42 | 0.51 | 0.70 - 2.87 | 0.98 | 0.33 |
|  | Bura | 1.00 |  |  |  |  |
| Constant |  | 0.40 | 0.14 | 0.20 - 0.82 | -2.53 | 0.01 |
| *Random effect* |  |  |  |  |  |  |
| Household ID |  | 1.22 | 0.65 | 0.43 - 3.44 |  |  |

Log Likelihood = -303.59; number of records = 607; number of groups = 302

The results given in Table 2 suggest that being a female was associated with higher odds of exposure to the pathogen that being a female. Similarly, pastoralists had significantly greater odds of exposure than famers and students. Ijara had significantly higher *Leptospira* spp. seroprevalence compared to all the other three sites (Bura, Sangailu and Hola).

Household ID, fitted as a random effects variable, was significant in the model (⎕^2^ = 8.24, p = 0.00). The residual intra-class correlation coefficient associated with this effect was 0.27 (95% CI: 0.12 – 0.51).

## Discussion

This study investigated the seroprevalence of *Leptospira* spp. in people who lived in pastoral (Garissa) and agropastoral (Tana River) counties, eastern Kenya. Leptospirosis is a neglected zoonotic disease whose distribution in the country is poorly known. It has also not been included in the list of diseases that should be screened for while investigating causes of febrile illnesses in humans. Our findings – which suggest a high seroprevalence of the disease in the study areas, more so in pastoral areas – show that awareness on the disease and its risk factors should be enhanced.

Both study regions used (Garissa and Tana River counties) lie on the either side of Tana River, a major river in Kenya that emanates from the slopes of Mt Kenya and terminates in the Indian Ocean. Their climatic conditions and vegetation cover provide good environmental conditions that support a wide range of wild animals. Livestock and wild animals act as reservoirs of *Leptospira* [16–18] and it is likely that silent transmissions of *Leptospira* spp. occur between livestock and wildlife since they share common grazing and watering resources.

One of the main but unexpected finding of the study was that *Leptospira* seroprevalence was significantly higher in the pastoral (Garissa) areas compared to agropastoral or irrigated areas in Tana River County. Outbreaks of leptospirosis is often precipitated by flooding and it was thought that irrigated areas would have higher seroprevalence. One possible explanation for this finding is that most of the people sampled in Garissa, especially those that lived miles away from River Tana, obtained water for domestic use from open water pans that had been built to trap rain water. These pans however served as common watering points for livestock and wildlife. They wade into the waters while drinking; they often urinate in and around these pans during or after taking a drink. People sampled in Tana River could access flowing water supplied to them via irrigation canals.

Water fetched from water pans in the pastoral areas could be boiled before use to reduce the risk of exposure to *Leptospira* spp. However, there would still be multiple opportunities for contact, especially for women and children who play a major role in fetching the water, or for the young men who took animals for watering in these pans. Our results showed that female gender, engaging in livestock related activities were associated with *Leptospira* seropositivity. We observed that age group of 21-30 were more likely to be seropositive. Higher infection rates in this age group corroborates findings in the majority of leptospirosis studies around the world, given the multiple but high risk responsibilities that people in this age group are assigned to (e.g. fetching water, taking care of animals, increased outdoor activities and recreational exposures, e.g. swimming). Seropositivity rates are lower in older age groups as risky exposure behaviours reduce [2,15].

Other gender-related activities that would force women in pastoral communities to come into direct contact with potentially contaminated water include washing, cleaning utensils, and cleaning their houses. This further shows that unlike other diseases where men often suffer higher risk of exposure given their outdoor occupations, *Leptospira* exposure may be more important in women than men in pastoral areas. Similar findings have been made in a study that used National health survey data in Chile [24].

Of importance to future *Leptospira* cross sectional studies in the region is the documentation of significant variation in the number of cases between households. An intra-cluster correlation coefficient of 0.27 (95% CI: 0.12 – 0.51) was estimated from the study. This demonstrates that households had significant differences on the extent to which they were exposed to the hazard. It therefore demonstrates that there is room to improve the management of *Leptospira* exposure in these communities for instance through behaviour change communication. The study had a few limitations though. It measured antibodies for *Leptospira* to identify evidence of prior infection. However, many *Leptospira* infections are subclinical. The development of clinical disease is dependent on multiple factors including pathogenicity of the serovars, individual’s immune status, comorbidities, and age. This may cause unnecessary alarm to the public health in terms of the high prevalence found.

We conclude that Leptospira is high in Garissa and Tana River counties. Females in pastoral areas had higher odds of exposure compared to males. Leptospirosis has not received sufficient attention despite the magnitude of the public health threat it poses and its negative economic impact on development and livelihoods in rural communities in low income countries. It is therefore important to gather data on the burden of leptospirosis to enable decision-makers to develop policy aimed at mitigating the effects based on reliable scientific evidence.

We recommend the inclusion of *Leptospira* surveillance in the disease surveillance and response (DSRU) platform being implemented by the Ministry of Health to determine the extent of the problem and its occurrence patterns. A ‘One Health’ approach to leptospirosis research and control should also be promoted to improve understanding of the epidemiology of the disease.

## Acknowledgements

We acknowledge the International Livestock Research Institute, University of Nairobi, Ministry of Health (Division of Disease Surveillance and response), Kenya Medical Research Institute, Kenyatta National Hospital, Director of Veterinary Services-Kenya and Kenya Field Epidemiology and Laboratory Training Program for their funding and/or collaboration in this study.

---

Figure 1: Flow chart showing the prevalence of Leptospirosis in Garissa and Tana River Counties, Kenya, December 2013-February 2015,

